# Ultrasound imaging links soleus muscle neuromechanics and energetics during human walking with elastic ankle exoskeletons

**DOI:** 10.1101/2020.01.20.909259

**Authors:** R.W. Nuckols, T.J.M Dick, O.N. Beck, G.S. Sawicki

**Author notes:** Corresponding authors: R.W. Nuckols and G.S. Sawicki.

## Abstract

Unpowered exoskeletons with springs in parallel to human plantar flexor muscle-tendons can reduce the metabolic cost of walking. We used ultrasound imaging to look ‘under the skin’ and measure how exoskeleton stiffness alters soleus muscle contractile dynamics and shapes the user’s metabolic rate during walking. Eleven participants (4F, 7M; age: 27.7 ± 3.3 years) walked on a treadmill at 1.25 m s^-1^ and 0% grade with elastic ankle exoskeletons (rotational stiffness: 0-250 Nm rad^-1^) in one training and two testing days. Metabolic savings were maximized (4.2%) at a stiffness of 50 Nm rad^-1^. As exoskeleton stiffness increased, the soleus muscle operated at longer lengths and improved economy (force/activation) during early stance, but this benefit was offset by faster shortening velocity and poorer economy in late stance. Changes in soleus activation rate correlated with changes in users’ metabolic rate (*p* = 0.038, R^2^ = 0.44), highlighting a crucial link between muscle neuromechanics and exoskeleton performance; perhaps informing future ‘muscle-in-the loop’ exoskeleton controllers designed to steer contractile dynamics toward more economical force production.

## INTRODUCTION

The plantar flexors serve an essential role in human walking by providing more than 50% of the leg’s total positive mechanical energy^1^ and up to 60% of the mechanical power output for redirecting the body’s center of mass during push-off ^2^. The importance of the ankle plantar flexors in legged locomotion is linked to their morphology. Muscle-tendons (MTs) with short pennate muscle fibers and long compliant in-series tendons are well suited for economical locomotion because the elastic tendons store and return of mechanical energy over each step^3-5^. During steady-state walking, the interaction of the plantar flexors and Achilles tendon is tuned. Throughout early stance, 0% (heel strike) to 40% stride, plantar flexor muscle fascicles generate force isometrically while the Achilles tendon stretches and stores mechanical energy^6^. In late stance, 40% to 60% (toe-off) stride, the plantar flexors shorten and the tendon rapidly recoils, providing a burst of positive mechanical power^1,4,7^. This coordinated MT interaction permits the plantar flexor muscle fascicles to operate over a narrow region of their force-length (F-L) curve and remain at slow shortening velocities which are favorable contractile conditions for economical force production^7-12^. Any disruption to this ‘catapult’ like mechanism may decrease the ankle’s efficient mechanical power production and worsen walking efficiency.

Exoskeletons are a class of wearable devices that often act in parallel with human MTs to restore or augment human movement. An increasing number of studies^13^ are establishing that both tethered^14-18^ and portable^19-21^ lower-limb exoskeletons can deliver mechanical power to the body to reduce metabolic demand during walking in young healthy individuals^14-22^, individuals post-stroke^23^, and older adults^24,25^. Recently, our group has demonstrated that exoskeletons need not deliver net external mechanical power to the body to reduce the metabolic rate of walking^26^. We showed that elastic exoskeletons that place a tension spring in parallel with the human plantar flexor-Achilles tendon complex can reduce the net metabolic rate of walking by an average of 7.2%^26^ versus not using a device. The relationship between the user’s net metabolic rate and the rotational stiffness of the exoskeleton is bowl-shaped, with a ‘sweet-spot’ that maximizes metabolic benefit where exoskeleton stiffness is not too compliant, is not too stiff, but just right.

Experimental evidence indicates that when exoskeletons are too stiff, a metabolic penalty arises from a series of compensatory changes in mechanics at the knee and hip as well as increased activation of the plantar flexor’s antagonist muscle, the tibialis anterior^26^. In addition to compensatory mechanisms at more proximal joints, local changes in ankle plantar flexor muscle dynamics may also help explain why increasing exoskeleton stiffness does not continue to improve metabolic rate during walking. Although increased exoskeleton stiffness reduces force requirements on the plantar flexors, the tuned MT interaction may become disrupted and result in poorer muscle economy (force/activation) and increased metabolic rate. This disruption of MT interaction which elicits less economical muscle dynamics is predicted by models and simulations of hopping^27,28^, walking^29-31^ and running^32^ and has been demonstrated experimentally in both human hopping^33^ and in isolated animal muscle experiments that use a biorobotic interface to simulate exoskeleton interaction dynamics^34^. During human hopping, an exoskeleton providing stiffness in parallel to the ankle plantar flexors reduced muscle force and net metabolic rate, but increased soleus muscle positive work due to greater fascicle excursions^33^. In a follow-up modelling study using the same hopping dataset, ankle exoskeleton torque reduced soleus fascicle operating lengths and increased fascicle velocities ^27^, which both acted to limit the estimated metabolic savings from using the device. Musculoskeletal simulations of walking with a unilateral elastic exoskeleton have demonstrated that applying ankle exoskeleton torque to the ankle-joint reduces muscle force requirements, which leads to decreased tendon stretch and increased fascicle strain^17^. This trend is also supported by a recent study that used ultrasound imaging to track the junction between the medial gastrocnemius and Achilles tendon during walking with a dynamic orthosis^35^. Together, these studies demonstrate that exoskeletons may disrupt the normal plantar flexor MT dynamics by limiting the ability of series-elastic tissues to keep plantar flexor muscles operating with contractile dynamics favorable for economical force production^30^.

Rather than relying on model-based estimates, we sought to use ultrasound imaging techniques to directly examine whether increasing ankle exoskeleton stiffness disrupts plantar flexor muscle-tendon dynamics during walking (Fig. 1). To date, no study has linked *in vivo* measurements of muscle dynamics to a user’s net metabolic rate during walking with exoskeletons. We used B-mode ultrasound imaging to measure soleus muscle fascicle contractile dynamics *in vivo* during exoskeleton assisted walking across a range of ankle exoskeleton rotational stiffnesses (k_exo_ = 0, 50, 100, 150, 250 Nm rad^-1^). We predicted that stiffer ankle exoskeletons would disrupt the normally tuned catapult behavior of the ankle plantar flexors. Specifically, we hypothesized that increasing exoskeleton stiffness would (1) reduce soleus muscle fascicle force, (2) reduce soleus activation, (3) increase soleus fascicle length and (4) shortening velocity, and (5) alter soleus force per unit activation (*i.e.*, muscle economy or force production capacity). Taken together, we posit that ankle exoskeleton assistance elicits a trade-off between reduced soleus force on one hand and a shift to contractile conditions that are less economical for force production on the other. We suggest that this balance explains the metabolic ‘sweet-spot’ at intermediate elastic ankle exoskeletons stiffness during walking.

**Figure 1:**
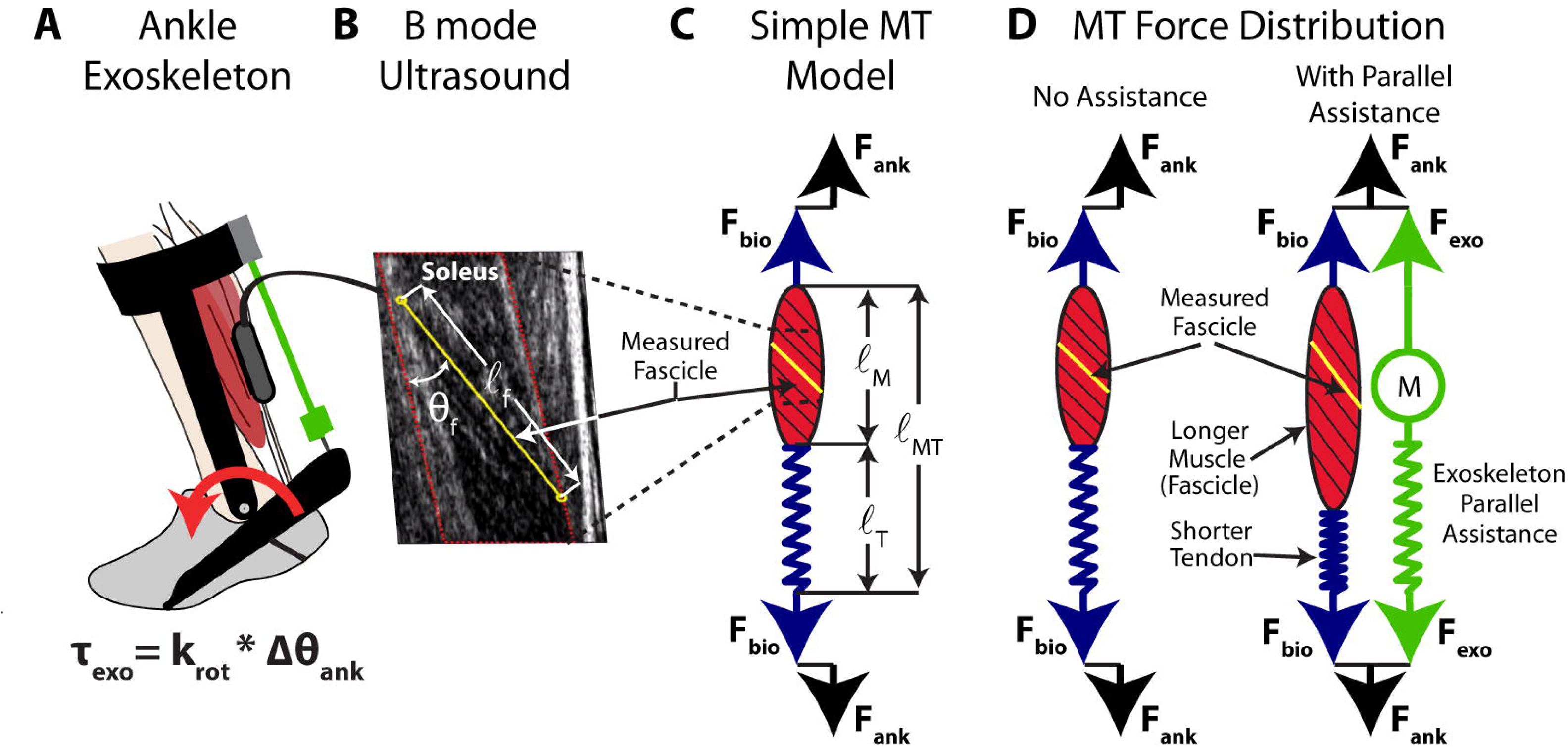
Conceptual model of an ankle exoskeleton providing parallel assistance to a plantar flexor muscle-tendon. **(A)** Schematic of an ankle exoskeleton that applies a plantar flexor torque about the human ankle joint. In this study, the torque behaved like a rotational spring (τ_exo_ = k_rot_ * Δθ_ank_) but conceptually could be any torque resulting from an exoskeleton force acting in parallel with a biological muscle-tendon. **(B)** We recorded B-mode ultrasound data from a probe attached to the shank over the soleus. We digitized images using automated tracking software to determine time-varying soleus fascicle lengths (*ℓ*_f_) and pennation angles (θ_f_). **(C)** In our simplified conceptual model, the muscle-tendon (MT) is comprised of a muscle (M) and tendon (T) (lengths = *ℓ*_mt_, *ℓ*_m_, and *ℓ*_t_ respectively) acting in series to produce a biological force (F_bio_) which generates a torque about the ankle through a moment arm (not shown for simplicity). **(D)** Without exoskeleton assistance (F_exo_ = 0), the entirety of the ankle force (F_ank_) is directed through the biological structure (F_bio_ = F_ank_). With exoskeleton assistance in parallel, a portion of the total ankle force is shunted to the exoskeleton resulting in a decrease in the biological force requirements (F_bio_ = F_ank_ - F_exo_). As a result of decreased biological force and because the tendon is a passive element, we expect the tendon strain to decrease by Δ*ℓ*_T_ = ΔF_bio_*(tendon stiffness). Finally, if length changes of the MT do not change much, we expect the muscle fascicles to be longer when exoskeleton parallel assistance is applied. In this study, we tested this hypothesis by using ultrasound to directly measure soleus fascicle length during walking with elastic ankle exoskeletons over a broad range of stiffnesses.

## RESULTS

To highlight the effect of exoskeleton stiffness on biomechanics and walking economy, the results focus on the effect of two exoskeleton stiffnesses with respect to the 0 Nm rad^-1^ (no assistance) condition. The 50 Nm rad^-1^ condition details what occurs with a stiffness that is low but results in the metabolic minimum. The 250 Nm rad^-1^ condition details what occurs at a perceptually very stiff condition which also resulted the metabolic maximum. Response for other conditions are reported in figures and Table 1.

**Table 1:**
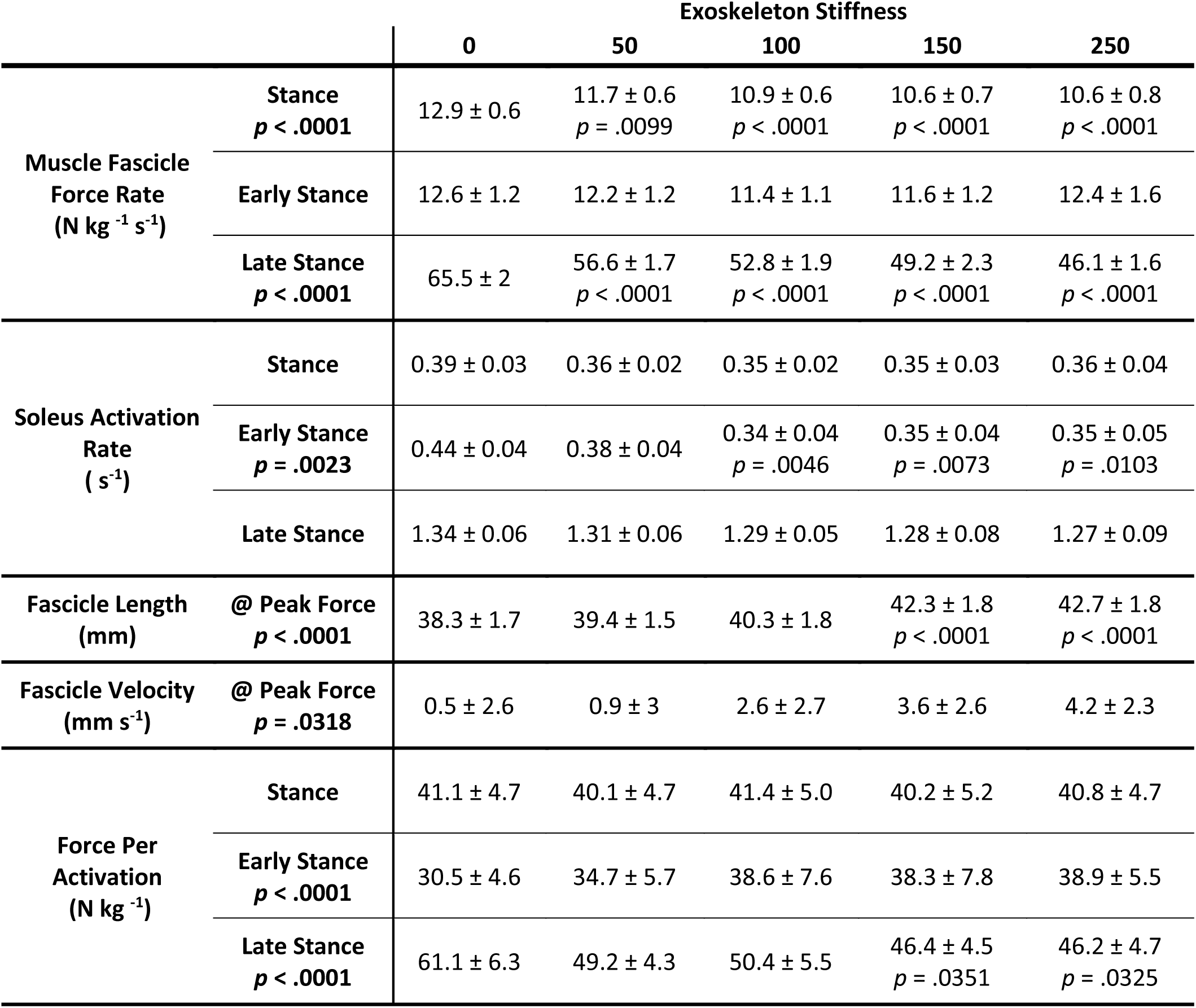
Muscle level changes due to exoskeleton assistance. Values are mean ± SEM across N=10 participants. [main effect: stiffness]. Post-Hoc test (THSD) on determine significant difference from 0 Nm rad^-1^. Early stance is defined as 0-40% of the gait cycle and late stance is 40-60%. (Note: The rate represents an average moment/activation per unit time and is not a measure of how rapidly the moment/activation is generated)

### Metabolic cost

We found a significant relationship between net metabolic rate and the square of exoskeleton stiffness (mixed-model ANOVA; k^2^_exo_; *p* = 0.022; Metabolic Rate 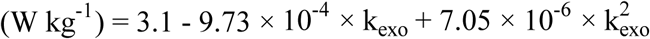). Elastic ankle exoskeletons providing 50 Nm rad^-1^ of rotational stiffness decreased net metabolic rate during walking at 1.25 m s^-1^ by 4.2% with respect to the 0 Nm rad^-1^ condition (CI:<0.4%, 7.8%>; two tailed paired t-test; *p* = 0.032) (Fig. 2A). Metabolic demand at 250 Nm rad^-1^ was 4.7% higher than 0 Nm rad^-1^.

**Figure 2:**
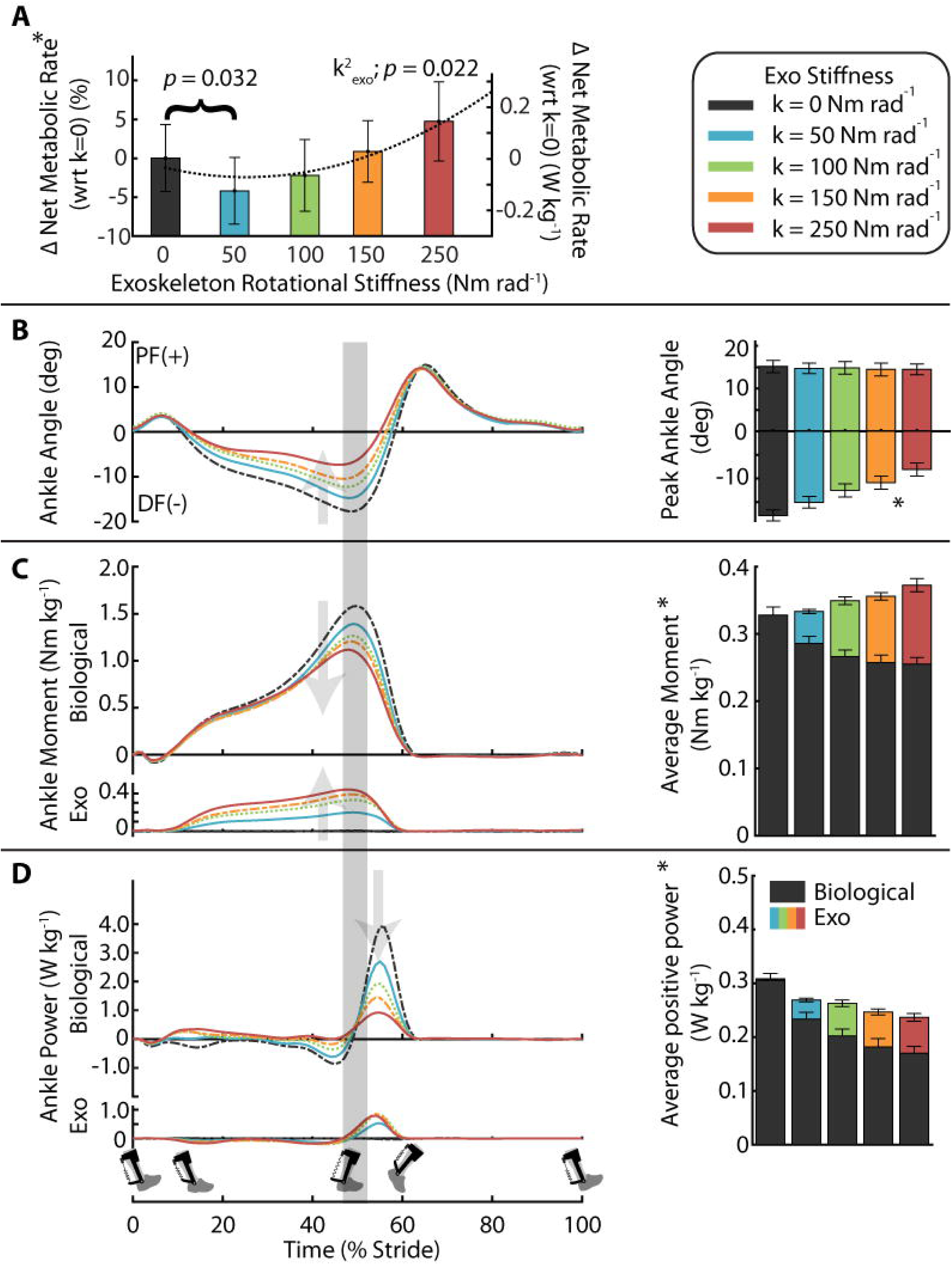
Effect of elastic ankle exoskeleton on metabolic demand and ankle joint dynamics. **(A)** Percent difference in metabolic rate for each stiffness relative to the zero-stiffness condition. Metabolic rate was minimized at 50 Nm rad^-1^ (ANOVA; k^2^_exo_; *p* = 0.022). **(B)** Time series of ankle joint plantar flexion (PF) and dorsiflexion (DF) angle averaged over participants. Peak ankle dorsiflexion angle was reduced with increased ankle exoskeleton stiffness. **(C)** Mass normalized biological moment (top) and exoskeleton torque (bottom) for each stiffness. Stacked bar charts represent the average biological (black-lower) and exoskeleton (colors-upper) contribution to total ankle moment for the stride. Increasing exoskeleton stiffness resulted in increased exoskeleton torque and decreased biological moment. **(D)** Time series of 6DOF biological (top) and exoskeleton (bottom) power for each stiffness. Stacked charts represent average biological (black, lower) and exoskeleton (colors, upper) power over the stride. Exoskeleton power remained fairly constant at high stiffnesses while biological power decreased with increased stiffness. Region of peak force highlighted in time series data. All bars graphs show mean ± SEM across N=11 participants. [main effect: stiffness, \**p* < 0.0001 (BIO, EXO, and ANK)]

### Spatio-Temporal Parameters

Increasing ankle exoskeleton stiffness reduced stride time (ANOVA; *p* = 0.0203) and stance time (ANOVA; *p* = 0.0012) but did not significantly affect swing time (ANOVA; *p* = 0.6454) or stance to swing ratio (ANOVA; *p* = 0.1559). Stride time and stance time decreased by ∼0.04 seconds (4%) at the stiffest condition compared to the 0 Nm rad^-1^ condition. Stance to swing ratio was ∼0.63 across all exoskeleton stiffness conditions.

### Ankle joint dynamics

Peak ankle dorsiflexion angle decreased as exoskeleton stiffness increased (ANOVA; *p* < 0.0001) (Fig. 2B). Relative to 0 Nm rad^-1^, peak ankle dorsiflexion decreased from 17.9 ± 1.2 deg. (mean ± SEM) to 15.0 ± 1.3 deg. for 50 Nm rad^-1^ and to 8.1 ± 1.4 deg. for 250 Nm rad^-1^. Increasing exoskeleton stiffness increased exoskeleton torque (ANOVA; *p* < 0.0001) and decreased biological ankle moment (ANOVA; *p* < 0.0001) (Fig. 2C). At 50 Nm rad^-1^, the exoskeleton provided a peak torque of 0.20 ± 0.01 Nm kg^-1^ and decreased biological ankle moment by 12% from 1.59 ± 0.04 Nm kg^-1^ to 1.41 ± 0.03 Nm kg^-1^. At 250 Nm rad^-1^, exoskeleton torque reached a peak of 0.47 ± 0.03 Nm kg^-1^ and reduced biological ankle moment by 28% to 1.14 ± 0.04 Nm kg^-1^. Increasing exoskeleton stiffness increased exoskeleton mechanical power (ANOVA; *p* < 0.0001) and reduced biological ankle mechanical power (ANOVA; *p* < 0.0001) (Fig. 2D). At 50 Nm rad^-1^, exoskeleton positive mechanical power was 0.036 ± 0.003 W kg^-1^ and average biological positive mechanical power decreased from 0.305 ± 0.013 W kg^-1^ to 0.233 ± 0.012 W kg^-1^. At 250 Nm rad^-1^, exoskeleton positive mechanical power was 0.066 ± 0.007 W kg^-1^ and average biological positive mechanical power decreased to 0.170 ± 0.012 W kg^-1^.

### Soleus muscle dynamics

Soleus muscle force decreased with increasing exoskeleton stiffness (Fig. 3A, Table 1). Peak force was greatest at the 0 Nm rad^-1^ condition, 21.88 ± 0.72 N kg^-1^, and minimized at the 250 Nm rad^-1^ condition, 15.18 ± 0.67 N kg^-1^. Reduced soleus force was reflected by decreased average force rate over the stance phase (0-60% stride) (ANOVA; *p* < 0.0001) (Fig. 3A) (Note: The rate represents an average moment per unit time and is not a measure of how rapidly the moment is generated). Compared to 0 Nm rad^-1^ condition, average soleus force rate of stance reduced by 12% and 18% for 50 and 250 Nm rad^-1^ respectively (Fig. 3Ai). There was no reduction in force rate of early stance (0-40% stride) (ANOVA; *p* = 0.38) (Fig. 3Aii). The change in average force rate of stance was primarily from the decrease in the average force rate of late stance (40-60% stride) (ANOVA; *p* < 0.0001) which, compared to 0 Nm rad^-1^, was reduced by 14% and 29% for 50 and 250 Nm rad^-1^ respectively (Fig. 3Aiii).

**Figure 3:**
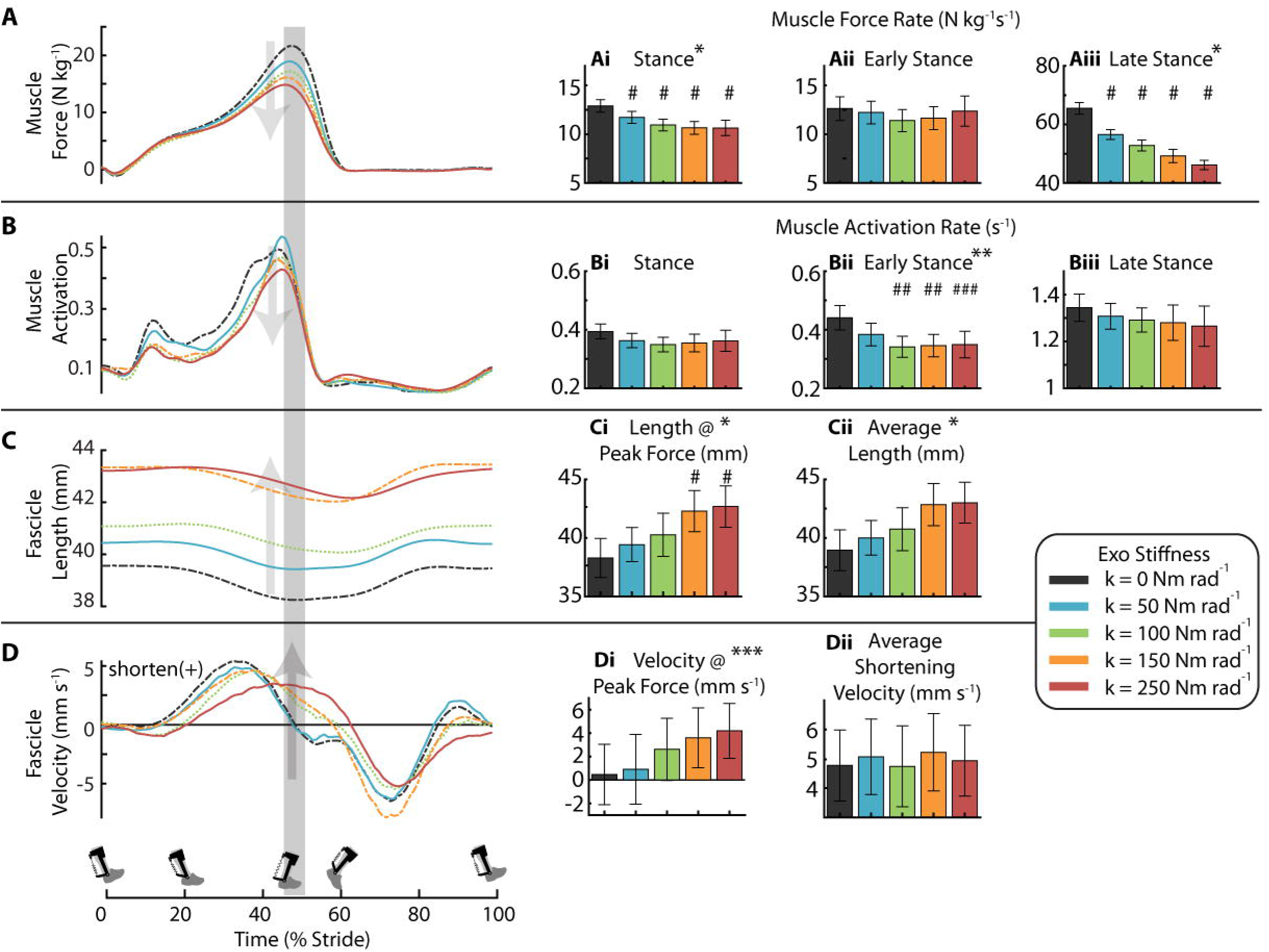
Effect of elastic ankle exoskeleton stiffness on soleus muscle-tendon dynamics. **(A)** Soleus force for each exoskeleton stiffness averaged over participants. Region of peak force highlighted in time series data. Bar charts represent the soleus muscle force rate for **(Ai)** stance, **(Aii)** early stance (0-40% stride), and **(Aiii)** late stance (40-60% stride). Increasing exoskeleton stiffness resulted in decreased soleus force rate for stance and late stance. **(B)** Time series of soleus muscle activation where amplitude is normalized to the peak activation across all stiffnesses for each participant. Bar charts are soleus muscle activation rate in **(Bi)** stance, **(Bii)** early stance, and **(Biii)** late stance. Soleus activation rate decreased with increasing exoskeleton assistance in early stance. **(C)** Time series of soleus fascicle length and bar charts representing **(Ci)** length at peak fascicle force and **(Cii)** average length during stance. Fascicle length increased with increasing exoskeleton stiffness. **(D)** Time series of soleus fascicle velocity measured and bar charts representing **(Di)** velocity at peak fascicle force and **(Dii)** average shortening velocity during stance. Fascicle shortening velocity at peak increases with stiffness. All bars graphs show mean ± SEM across N=10 participants. [main effect: stiffness, \**p* < 0.0001, \*\**p* = 0.0023, \*\*\**p* = 0.0318; THSD: ^#^*p* < 0.0001, ^##^*p* < 0.01, ^###^*p* < 0.05].

Soleus average activation rate, as measured by surface electromyography (EMG), decreased with increasing exoskeleton stiffness during early stance (ANOVA; *p* = 0.0023) (Fig. 3B and Table 1). The reduction in activation rate of early stance was 14% and 21% for the 50 and 250 Nm rad^-1^ condition compared to 0 Nm rad^-1^ (Fig. 3Bii). There was no change in activation rate of late stance (ANOVA *p* = 0.8765) (Fig. 3Biii).

The peak length of the soleus MT decreased with increasing exoskeleton stiffness (ANOVA; *p* < 0.0001). Compared to 0 Nm rad^-1^, MT peak length decreased from 308 ± 6.6 mm to 306 ± 6.9 mm in the 50 Nm rad^-1^ condition and to 301 ± 6.9 mm in the 250 Nm rad^-1^ condition (Supp Fig. 1C).

Ankle exoskeletons altered both soleus fascicle length and velocity dynamics. Stiffer exoskeletons increased fascicle average length during all of stance (ANOVA; *p* < 0.0001) and during the time of peak force (ANOVA; *p* < 0.0001) (Fig. 3C, 4A, Table 1). Compared to fascicle length at 0 Nm rad^-1^ (38.3 ± mm), fascicle length at the time of peak force increased by 2.9% and 11.4% in the 50 and 250 Nm rad^-1^ conditions respectively (Fig. 3Ci, 4A); likely shifting the respective fascicles toward their optimal length (*l*_*0*_) on the F-L curve (Fig. 4C).

**Figure 4:**
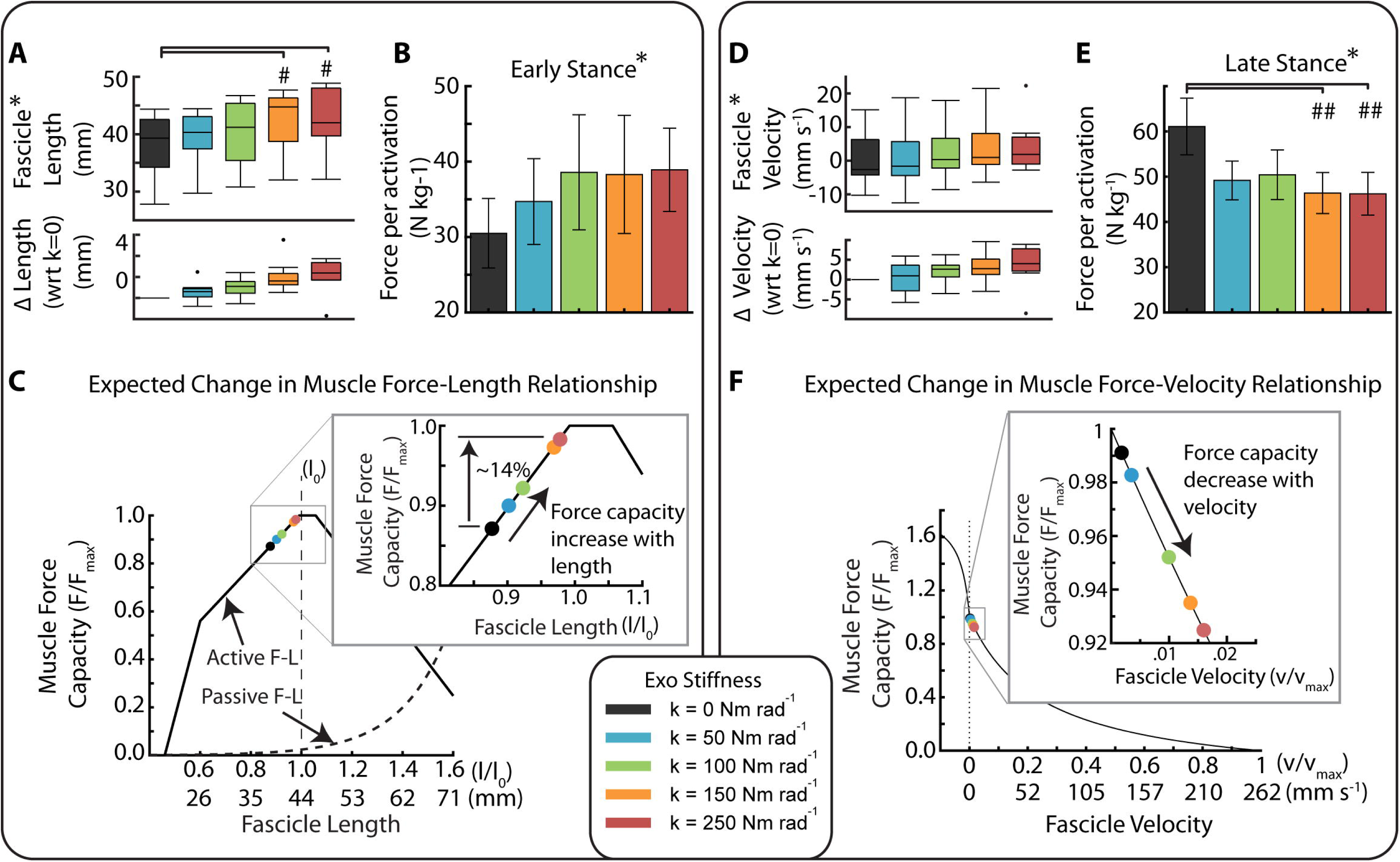
Trade-off in force-length-velocity behavior links users’ soleus fascicle dynamics and economy of force production. **(A)** (Top) Soleus fascicle length at peak fascicle force for each stiffness. (Bottom) Difference in soleus fascicle length at peak force relative to the zero stiffness conditions. The two high stiffness conditions (orange and red) were significantly longer than 0 Nm rad^-1^ (ANOVA \**p* < 0.0001; THSD ^#^*p* < 0.0001). **(B)** Average muscle force per activation (*i.e.*, muscle economy) during early stance (0-40%) of the gait cycle. Muscle economy increased (\**p*<0.0448) with increased exoskeleton stiffness as might be expected from improved operating point on the F-L curve during early stance. **(C)** Fascicle operating length at peak force relative to estimated muscle force-length (F-L) curve with optimal length *l*_*0*_ = 43.6 mm and inset of a narrowed scale. Data indicates a rightward shift of the F-L curve with increased exoskeleton stiffness and increased force production capacity. **(D)** (Top) Soleus fascicle velocity at peak fascicle force for each stiffness (\**p*=0.032). (Bottom) Difference in soleus fascicle velocity at peak force relative to the zero stiffness conditions. **(E)** Average muscle force per activation during late stance (40-60%) of the gait cycle. Muscle economy decreased with increased exoskeleton stiffness as might be expected from a shift in operating point on the F-V curve during late stance. The two high stiffness conditions (orange and red) were significantly different than 0 Nm rad^-1^ (ANOVA \**p* = 0.0258; THSD ^##^*p* = 0.0351 & 0.0325). **(F)** Fascicle operating point at peak force on estimated muscle force-velocity (F-V) curve assuming v_max_ of 262 mm s^-1^. Shift in muscle velocity to the right with increased stiffness shown in inset due to narrow range of shortening velocities near v = 0 mm s^-1^. All bars graphs show mean ± SEM across N=10 participants.

Soleus fascicles shortened faster during stance with increasing exoskeleton stiffness, and this was reflected by increased fascicle velocity at the time of peak force (ANOVA; *p* = 0.0318) (Fig. 3D, 4D, Table 1). Compared to the fascicle velocity at the time of peak force for 0 Nm rad^-1^ (0.47 ± 2.6 mm s^-1^), fascicle shortening velocity increased by 0.4 ± 1.2 mm s^-1^ and 3.7 ± 1.6 mm s^-1^ in the 50 and 250 Nm rad^-1^ conditions, respectively (Fig. 3Di, 4D), shifting the soleus fascicles towards their maximum shortening velocity (Fig. 4F). No condition was significantly different from 0 Nm rad^-1^ (THSD: *p* > 0.05).

Ankle exoskeletons had no effect on soleus fascicle pennation angle during the stance phase. Neither the pennation angle at the time of peak force (26.6 ± 0.8 deg; ANOVA; *p =* 0.1007) nor the average pennation angle over stance (25.8 ± 0.8deg; ANOVA; *p=*0.2301) changed across exoskeleton stiffness conditions (Supp Fig. 1D).

Exoskeletons affected the economy of soleus force production in a phase dependent manner during stance; likely improving it during early stance and worsening it during late stance. Soleus muscle economy increased in early stance (ANOVA; *p* = 0.0448) (Fig. 4B, Table 1), increasing by 14% and 27% for 50 and 250 Nm rad^-1^ respectively compared to 0 Nm rad^-1^. Soleus muscle economy decreased in late stance (ANOVA; *p* = 0.0258) (Fig. 4E), decreasing by 20% and 24% at 50 and 250 Nm rad^-1^ compared to 0 Nm rad^-1^.

### Association between soleus neuromechanics and whole-body net metabolic rate

We found a positive correlation (LLSR: *p* < 0.038, R^2^ = 0.44) (Table 2, Fig. 5) between the change in average soleus activation rate of stance (*i.e.*, Δ = with respect to the zero exoskeleton stiffness condition) and Δ in whole-body net metabolic rate. In addition, we found a significant positive correlation (LLSR: *p* < 0.016, R^2^ = 0.42) (Table 2) between Δ soleus force rate in early stance phase and Δ whole-body net metabolic rate. Finally, we found a negative correlation between Δ soleus force (LLSR: *p =* 0.0018, R^2^ = 0.53) and Δ soleus force rate (LLSR: *p =* 0.0005, R^2^ = 0.59) (Table 2) during late stance and Δ whole-body net metabolic rate.

**Table 2.** The relationships between the change in net metabolic rate (W kg^-1^) (y) and changes in soleus muscle neuromechanics (x) during walking with exoskeletons over a range of stiffness values, k_exo_. For each metric, Δ represents the difference from the k_exo_=0 condition (*i.e.*, the effect of increasing exoskeleton stiffness) with positive values indicating an increase. We used a within-participant linear regression analysis to determine whether relationships were significant (*p* < 0.05), and for those that were, we report the line of best fit to the data and the R^2^ value of the fit. **<u>Bold underline</u>** indicates a significant relationship between the change in a given neuromechanical variable, x and the change in whole-body net metabolic rate, y.

| Gait Cycle Phase<br>(% Stride) | Total Stance (TS)<br>0-60% | Early Stance (ES)<br>0-40% | Late Stance (LS)<br>40-60% |
| --- | --- | --- | --- |
| $\Delta$ Muscle Positive Power [Fig. 2D]<br>($\text{J kg}^{-1} \text{s}^{-1}$) | $p=0.97$ | ---- | ---- |
| $\Delta$ Muscle Force<br>($\text{N kg}^{-1} \text{s}^{-1}$) | $p=0.45$ | $p=0.09$ | **<u>$y = -0.27 - 0.07x$</u>**<br>**<u>$p=0.0005, R^2=0.59$</u>** |
| $\Delta$ Muscle Force Rate [Fig. 3A]<br>($\text{N kg}^{-1} \text{s}^{-1}$) | $p=0.37$ | **<u>$y = 0.01 + 0.03x$</u>**<br>**<u>$p=0.016, R^2=0.42$</u>** | **<u>$y = -0.26 - 0.02x$</u>**<br>**<u>$p=0.0018, R^2=0.53$</u>** |
| $\Delta$ Muscle Activation<br>(unitless) | $p=0.41$ | $p=0.41$ | $p=0.64$ |
| $\Delta$ Muscle Activation Rate [Fig. 3B]<br>($\text{s}^{-1}$) | **<u>$y = 0.03 + 1.27x$</u>**<br>**<u>$p=0.038, R^2=0.44$</u>** | $p=0.08$ | $p=0.57$ |
| $\Delta$ Force / $\Delta$ Muscle Activation [Fig. 4B, E]<br>($\text{N kg}^{-1}$) | $p=0.64$ | $p=0.91$ | $p=0.63$ |

**Figure 5:**
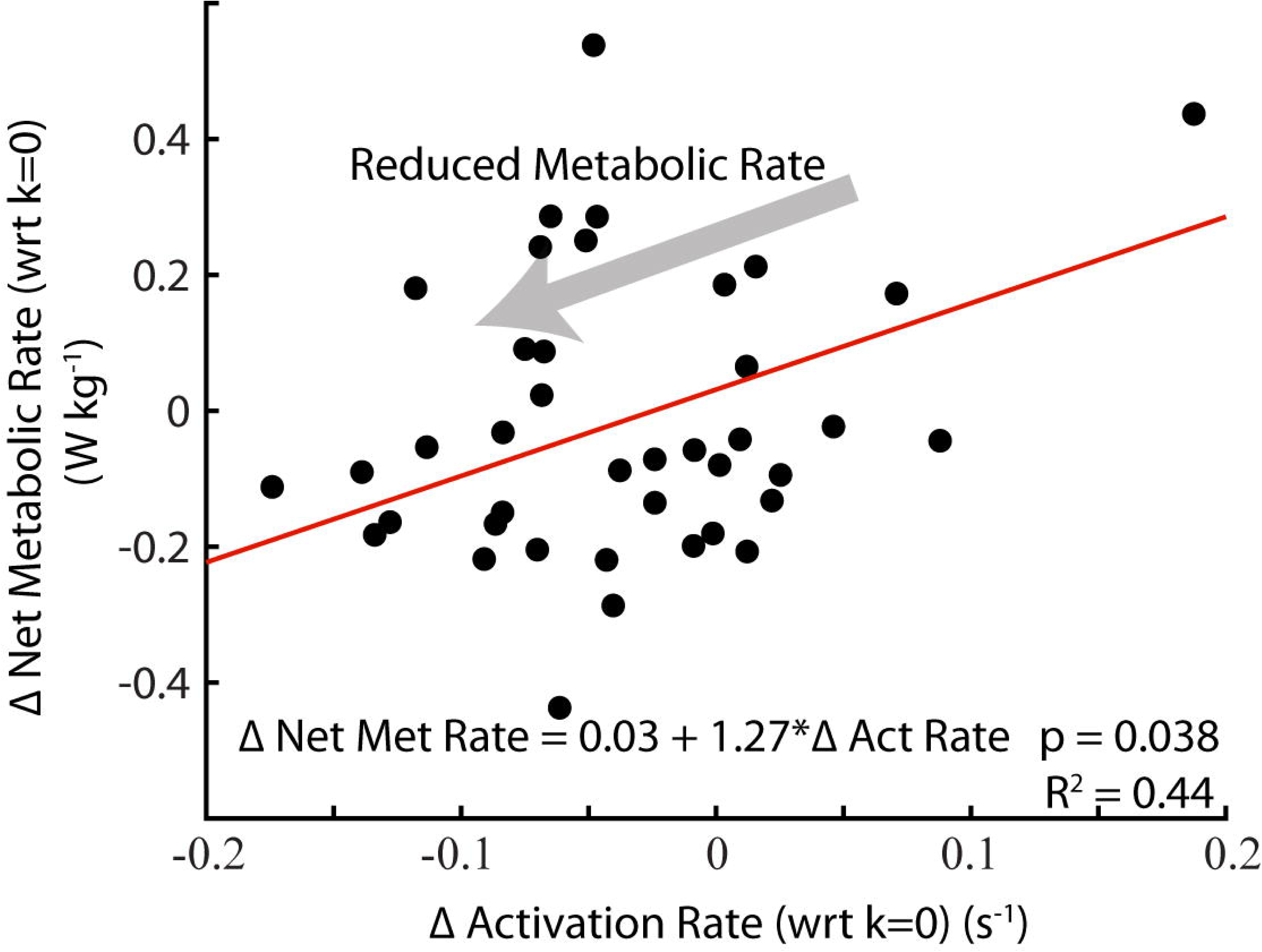
Relationship between rate of muscle activation and metabolic rate. Change net metabolic rate (W kg ^-1^) plotted against change in soleus activation rate (unitless s^-1^) with respect to no assistance (k = 0 Nm rad^-1^). Changes in soleus activation rate were significantly correlated with changes in users’ net metabolic rate (*p* = 0.038; R^2^ = 0.44.

## DISCUSSION

This study’s overarching hypothesis was that despite reducing plantar flexor force requirements during walking, elastic ankle exoskeletons would detune the catapult-like interaction within the plantar flexor MTs, affect the economy of plantar flexor force production, and yield a muscle-level metabolic change. As hypothesized, applying rotational stiffness to the ankle with elastic exoskeletons altered plantar flexor muscle dynamics. At a walking speed of 1.25 m s^-1^, increasing exoskeleton rotational stiffness resulted in decreased soleus force (Fig. 3A) and increased soleus fascicle length and shortening velocity at the time of peak force (Fig. 3C, D and Fig. 4A, D). These results provide the first empirical evidence to support predictions from previous modelling and simulation studies that used an *in silico* approach to examine changes in plantar flexor muscle-dynamics during walking with spring-loaded^30^ and actively powered^29^ ankle exoskeletons. Our approach provides an *in vivo* experimental framework that can be used to explore how exoskeletons may be used to steer (*i.e.*, direct) muscle fascicle dynamics to reduce users’ net metabolic rate^36^.

A muscle’s capacity to produce force is, in part, governed by the intrinsic force-length (F-L) and force-velocity (F-V) relationships^9-11,37-39^. In human walking, soleus fascicles normally operate on the ascending limb of the F-L curve^8^. While we did not characterize our participants’ soleus fascicle optimal lengths (*l*_*o*_), the average fascicle length during walking with zero exoskeleton stiffness (0 Nm rad^-1^) was ∼39 mm, which is up to 11% shorter than soleus optimal lengths reported in previous studies^40^. A study that mapped participants’ soleus length changes during walking at self-selected speed (mean = 1.14 ms^-1^) onto their empirically determined F-L curves found that humans operate at a nearly constant fascicle length of ∼0.9 *l*_*0*_ over early stance^8^. Assuming those in our study also operated at ∼0.9 *l*_*0*_, we estimate the soleus optimal length of 43.6 mm (39.2 mm/0.9) for our participants.

When operating on the ascending limb of the F-L curve, a shift to longer fascicle lengths should increase muscle force capacity, especially when considering the contribution of passive muscle forces at lengths greater than the optimal length^41^. Using equations for theoretical active and passive F-L relationships taken from Golapudi and Lin^42^ and our estimate of group average *l*_*0*_ of 43.6 mm, we calculated that the ∼4 mm increase in soleus fascicle length at 250 Nm rad^-1^ (Figs. 3C, 4A) would shift the muscle’s operating length onto the plateau of the F-L curve and therefore could increase soleus force capacity by as much as 12% compared to 0 Nm rad^-1^ (Fig. 4C). Thus, walking with increased ankle exoskeleton stiffness likely resulted in a length-dependent increase in soleus force capacity during early stance.

With respect to soleus fascicle shortening velocity, muscle force capacity is most sensitive to changes in velocity near the isometric region of the F-V curve^10^, and a shift to faster shortening velocities decreases force capacity^10^. In this study, for 0 Nm rad^-1^ the average fascicle velocity during stance (∼5 mm s^-1^) and velocity at the time of peak force (∼0.5 mm s^-1^) were considerably less than the reported maximum shortening velocity (*v*_*max*_) of the soleus (∼6\**l*_*o*_, 262 mm s^-1^)^30^ (Fig. 3D). These slow shortening velocities coincide with previous research suggesting that the soleus operates near isometrically during early stance in walking^7,8,12^. Using an equation from Alexander 1997^43^ and a *v*_*max*_ of 262 mm s^-1^, we calculated that the relatively small ∼4.0 mm s^-1^ increase in soleus fascicle shortening velocity at the time of peak force (Figs. 3D, 4D) for the 250 Nm rad^-1^ condition would reduce force capacity by 6.7% compared to 0 Nm rad^-1^ (Fig. 4F). Thus, walking with increased ankle exoskeleton stiffness likely resulted in a velocity-dependent decrease in soleus force capacity during late stance. The impact of increasing exoskeleton stiffness on muscle length and velocity dynamics presents an interesting trade-off with respect to the metabolic energy cost of soleus force production.

Inspired by a plethora of locomotion research linking changes in metabolic rate (*Ėmet*) to changes in rate of active muscle volume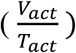 ^44-50^ 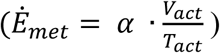, we examined the factors that influence the rate of active soleus muscle volume during walking with elastic ankle exoskeletons (see Supplementary Analysis (S1) for detailed analysis). We found evidence that elastic ankle exoskeletons influence whole-body net metabolic rate *Ėmet* primarily through their impact on soleus muscle activation rate (Fig. 3B) during the stance phase of walking (Table 2, Fig. 5). Our mechanistic framework (S1, Eqs. S1-S7) revealed that elastic exoskeletons modified soleus activation rate not just by modulating force demand (S1, Eqs. S3-S4; Fig. 3A), but also by modifying soleus length change dynamics (Figs. 3C,4A and Figs. 3D,4D) – shifting the muscle’s operating point on its F-L and F-V curves (Fig. 4C, F) subsequently impacting its force capacity (S1, Eqs. S5-S7; Fig. 4 B, E). These findings represent the first empirical evidence that show changes in local plantar flexor muscle dynamics well-explain the ‘bowl-shaped’ relationship between elastic ankle exoskeleton stiffness and user’s net metabolic rate ^26,51^.

The rate of soleus active muscle volume 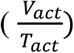 captures the changes in local muscle dynamics (*e.g.*, force, length, velocity, and ultimately activation) due to elastic ankle exoskeleton torque and can explain a significant portion (∼44%, (Table 2)) of the change in whole-body net metabolic rate with increasing exoskeleton stiffness. Although substantial, the rate of soleus active muscle volume explains less than 50% of the change in net metabolic rate across exoskeleton stiffness. Even though the soleus is presumably one of the body’s most metabolically active muscles during locomotion due to its relatively large force output^47-49,52,53^, and ankle exoskeletons likely alter muscle dynamics at the ankle more than knee or hip muscle dynamics^26,27,54^, there are other muscles^55^ and factors that contribute to whole-body metabolic energy expenditure^55^.

From a theoretical and purely mechanical perspective, users should be able to maximize the benefits of the exoskeleton by replacing the biological moment with an equivalent contribution from the exoskeleton. To do this, users would ‘shut off’ their plantar flexor muscles to minimize the biological moment/power requirements and take full advantage of exoskeleton moment/power. However, we found that the total (biological plus exoskeleton) net ankle moment increased with increasing exoskeleton stiffness (Fig. 2C), indicating that users did not equally reduce the forces in their plantar flexors (Fig. 3A) by the amount that would have been required to maintain a consistent total ankle moment. In addition to our results, many other studies^51,56-59^ also indicate that as exoskeleton assistance increases, humans do not ‘shut off’ their muscles and rely on exoskeleton assistance alone; instead a cascade of local and global neuromechanical compensations arise. For example, we observed decreased ankle dorsiflexion (Fig. 2B) and a plateau in exoskeleton torque across the highest stiffness values (Fig. 2C), along with increased tibialis anterior activity^51^ and knee flexion moment at terminal stance^51^. These compensations may functionally reduce injury potential, as studies have shown that the biological system may be resistant to increased strain on the muscles as a mechanism to prevent eccentric muscle damage^60-63^. We observed a ∼10% increased soleus fascicle strain from the 0 to the 250 Nm rad^-1^ exoskeleton condition (Figs. 4A,C), well below the strain levels that have been reported to be damaging (>1.25 *l*_*o*_)^63^. However, if users had not reduced their dorsiflexion in late stance as much as they did (Fig. 2B) (∼ equivalent to 7 mm MT excursion (see Supp Fig. 3C)) while maintaining the same torque profile, we predict that peak fascicle lengths would have approached the 25% > *l*_*0*_ level. Furthermore, the observed compensations in ankle kinematics (*e.g.*, increased plantar flexion angle) were similar to the kinematic compensations observed during a study of human hopping with elastic ankle exoskeletons^33^ where injury prevention may have also been a factor.

Another potential explanation for users not taking advantage of maximum exoskeleton stiffness could be based in an inability to reduce their soleus muscle activation in proportion to the level of assistance provided. Research has shown that joints (and limbs) tend to maintain constant stiffness during a given locomotor condition^37^ which may be accomplished through a combination of stretch sensors (*e.g.*, muscle spindle organs) and force sensors (*e.g.*, golgi tendon organs) that are responsible for maintaining a nearly constant ratio between the change in muscle force and length^64-66^. Because exoskeleton torque decreased soleus force and increased fascicle length, the apparent stiffness of the MT was likely markedly reduced. Therefore, a mismatch in sensory feedback from the force (golgi tendon organ) and length (spindle organ) related mechanoreceptors could be a possible explanation for why we do not observe reductions in activation proportional to the level of applied exoskeleton assistance. Limb-joint stiffness control may be further regulated through higher neural centers, but much of walking is automatic and unconsciously controlled though spinal-level central pattern generators, reflexes, cerebellar regulation, and the brainstem^67^. For this reason, exoskeleton users may not be able to easily turn off their plantar flexors, which would be required to further minimize whole-body net metabolic rate across a broad range of stiffnesses. Further work in both humans and non-human animals^34,68^ is necessary to understand the interaction between altered muscle fascicle dynamics and feedforward/feedback neural control mechanisms during walking with exoskeletal devices.

We acknowledge certain potential limitations. The sessions were split among a training day and two testing days and there is the potential for day to day variability. Our goal was to balance the conditions that could be studied with the fatigue of the individual. Because individuals were trained and healthy young adults with no muscular or neurological issues, we thought the tradeoff acceptable. In our analysis and interpretation of muscle fascicle dynamics, while we were able to measure the dynamics of the muscle using ultrasound, certain measurements were simplified or calculated based on common assumptions. We did not measure optimal fascicle length directly but estimated that the optimal fascicle length is ∼0.9\**l*_*0*_ ^8^. The Achilles Tendon moment arm is geometrically derived^69^ rather than capturing the full complexity^70,71^. Due to our inability to directly measure soleus force and power, we used joint torque and assumed a distribution among redundant plantar flexor muscles based on relative muscle cross-sectional areas^72^ and no antagonist co-contraction.

In terms of application, our results highlight the importance of the morphology of the MTs that a device interacts with, unpowered or powered. Exoskeleton induced changes to the F-L operating point may provide an even greater benefit in populations where MT dynamics are already disrupted. Given the non-linear curvature of muscle’s F-L relationship, a given shift in muscle length could induce a larger improvement in economy when starting from even shorter lengths. For example, Achilles tendon compliance increases with advanced aging^73^, stroke^74^ or paralysis^75^ and this likely shifts the plantar flexor muscles to shorter operating lengths and faster shortening velocities, conditions that increase metabolic rate^76,77^. Furthermore, in cases where the foot-ankle complex becomes stiffer (*e.g.*, diabetic neuropathy^78^), intervening by adding stiffness to the foot instead of the ankle joint may improve plantar flexor capacity by yielding slower, more economical shortening velocities. Finally, a device working in parallel with a stiffer MT than that of the ankle (*e.g.*, hip flexors or extensors) might reduce underlying muscle activation predominantly by altering force demand without influencing force capacity (*i.e.*, F-L, F-V operating points) due to a reduced decoupling between the muscle and joint.

Regarding the design and development of future exoskeletons, our results reaffirm that net power transfer from an exoskeleton is not necessary to reduce net metabolic rate during locomotion, even for young adults whose plantar flexor neuromechanics are near optimally tuned for economical force production^21,31^. This is primarily because net mechanical work from a device is not necessary to reduce the muscle activation rate of the user, a key factor driving metabolic savings from exoskeleton assistance (Table 2, Fig. 5). Since muscle activation rate is the metric that best captured the complex trade-off between changes in force demand and force capacity derived from exoskeleton assistance, it may be a useful foundation to build an updated roadmap for exoskeleton design. Typically, engineers and scientists have focused on building exoskeletons to reduce lower-limb joint moments and powers in order to reduce metabolic rate and is reflected in exoskeleton performance measurements (*e.g.*, performance index^79,80^; augmentation factor^19,59^) that primarily relate reductions in biological mechanical power and net metabolic power of the user. But our results indicate no relationship between changes in users’ metabolic rate and changes in mechanical power of the soleus muscle due to exoskeleton assistance (Table 2). In addition, other lines of evidence suggest that changes in biological mechanical power at the center of mass, joint-, or muscle-level^14,36,56^ are unable to explain how exoskeletons alter users metabolic rate. Taken together, these results motivate the need to incorporate muscle-level analyses into the design process for wearable devices intended to influence the dynamic function of musculoskeletal tissues.

The muscle-level links that we established between device function and user performance may provide guidance for achieving a more complete symbiosis between human and machine. Exoskeletons that can directly and seamlessly target muscle dynamics may provide even greater locomotion performance benefits than current devices, perhaps beyond improving walking economy. One approach, inspired in part by recent animal work where *in vivo* muscle lengths were used for closed-loop control of muscle force^81^, is to develop exoskeleton controllers that can sample muscle states in real-time and then steer (direct) them by updating device parameters in real-time^36^. This remains a formidable challenge in humans where direct measurements of underlying muscle dynamics are so far, too invasive. However, there is potential to use advanced tools from image processing and machine-learning to extract real-time muscle length and velocity samples from ultrasound images^82^, and then use them in combination with EMG or a musculoskeletal model^83,84^ as input to an exoskeleton controller. Access to muscle dynamics in real-time would enable application of state-of-the-art human in the loop optimization techniques^17,85,86^ to individual or groups of target muscles. The possibility of ‘muscle in the loop’ exoskeleton control strategies opens the door to devices capable of steering neuromechanical structure and function over both short and long timescales during locomotion^87,88^. The potential impact of wearable devices ranges from improving a runner’s metabolic economy during a marathon; to altering sensory feedback for optimal rehabilitation post-stroke; to increasing the stiffness of the Achilles tendon to counteract the effects of aging. By implementing wearable devices that are symbiotic and highly adaptable, we have the potential to have a transformational impact on augmenting healthy and restoring pathological movement.

## CONCLUSIONS

We used B-mode ultrasound imaging to measure *in vivo* soleus contraction dynamics and demonstrated that ankle exoskeleton rotational stiffnesses altered the normally tuned catapult behavior of the ankle plantar flexors. As stiffness increased, fascicle length and velocity increased and likely affected the muscle force production capacity as reflected in changes to the muscle force per activation calculation. Despite the general reduction in fascicle force, the altered contractile dynamics help explain the ‘bowl-shape’ where an intermediate stiffness rather than high stiffness results in maximum metabolic benefit. This work highlights the importance of linking exoskeleton-related changes in muscle dynamics to changes in users’ metabolic rate and has implications for the design, application, and future development of exoskeletons beyond unpowered devices acting at the ankle.

## METHODS

### Participants

Eleven healthy adults (4 female, 7 male, age: 27.7 ± 3.3 years, height: 1.75 ± 0.07 m, mass: 76.8 ± 8.2 kg) participated in the study. Participants had limited experience with the elastic ankle exoskeleton prior to the study. All participants signed an informed consent to participate in the study which was approved by the Institutional Review Board at The University of North Carolina at Chapel Hill. The methods were carried out in accordance with the IRB approved protocol.

### Ankle exoskeleton testbed

The exoskeleton end effector was a lightweight carbon fiber ankle foot orthosis that applied plantar flexor torque to the ankle. The device consisted of bilateral ankle exoskeletons driven by benchtop motors (Fig. 6A). The control system imposed a torque angle relationship to emulate an elastic device providing rotational stiffness in parallel with the ankle. We calculated desired torque,*τ*_*exo*_ based off a predefined rotational stiffness and the real-time ankle angle measured using an electrogoniometer mounted to the exoskeleton shank and foot sections (Eq. 1)

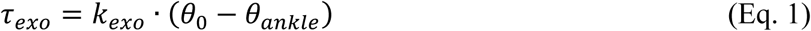

where *k*_*exo*_ was exoskeleton rotational stiffness, *θ*_0_ was the onset angle, and *θ*_*ankle*_ was the real-time ankle angle (Fig. 6C).

**Figure 6:**
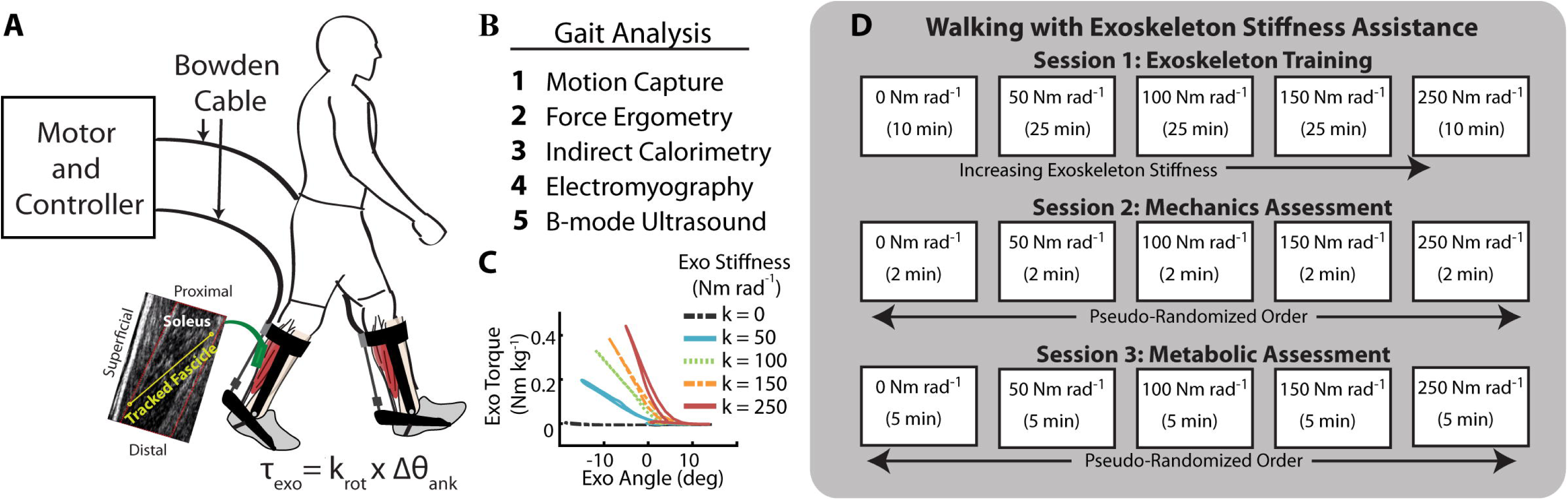
Elastic ankle exoskeleton test-bed and study protocol. **(A)** Representative setup of the exoskeleton testing platform for evaluating rotational stiffness plantar flexion assistance. The exoskeleton provided plantar flexion torque to the bilateral ankle exoskeletons through off-board motors. The controller emulated elastic rotational stiffness by imposing a torque angle relationship control law (τ_exo_ = k_rot_ * Δθ_ank_). **(B)** In addition to traditional gait analysis tools, we used B-mode ultrasound to track soleus fascicle length changes while exoskeleton assistance was applied. **(C)** Stiffness could be changed in software and representative torque-angle profiles at each stiffness are shown. **(D)** Protocol for the three sessions. The imposed waiting period between each testing session was 2-7 days to allow for learning and retention^89^ and reduce fatigue. Participants were trained in the first session on all stiffnesses at 1.25 m s^-1^. During the second session, we recorded joint mechanics, electromyography (EMG), and ultrasound data for 5 exoskeleton conditions (k_rot_ = 0, 50, 100, 150, 250 Nm rad^-1^). In the third session, we recorded steady state metabolic cost for the same conditions.

### Walking trials

Participants completed testing over three sessions where they walked at 1.25 m s^-1^ at five ankle exoskeleton rotational stiffness conditions (k_exo_ = 0, 50, 100, 150, 250 Nm rad^-1^). The order of the three testing sessions was (1) exoskeleton training, (2) gait mechanics, (3) steady-state metabolic energy consumption (Fig. 6D). The imposed waiting period between each testing session was 2-7 days to allow for learning and retention^89^.

1. **Training:** Previous work has demonstrated the importance of training on the acceptance of mechanical assistance in exoskeletons^80,90,91^. Based off the training time from other studies^26,80^ and the time reported in other studies as the time required to metabolically adapt^91^, each participant walked in the exoskeleton at 1.25 m s^-1^ for a total of 95 minutes over 5 training trials. The participants walked at each of intermediate stiffness conditions (k_exo_ = 50, 100, 150 Nm rad^-1^) for 25-minute trials and at the lowest and highest stiffness conditions (k_exo_ = 0, 250 Nm rad^-1^) for 10 minutes. We collected indirect calorimetry data for each of the trials to monitor changes in metabolic energy use over the course of the training session.
2. **Gait mechanics:** Participants walked for 2 minutes at each of the 5 exoskeleton stiffness trials in a randomized order. We instrumented participants with surface electromyography on the left shank (120Hz, Vicon, Oxford, UK), and motion capture markers on the lower limbs and pelvis. Due to space constraints, B-mode ultrasound was placed over the right soleus.
3. **Steady state metabolic energy expenditure:** We collected indirect calorimetry data for each walking trial. To allow participants’ metabolic rate to reach steady state, each walking trial lasted 5 minutes. The order of the steady state metabolic conditions was re-randomized.

### Metabolic energy consumption measurements

We calculated mass-specific net metabolic power (W kg^-1^) using a portable indirect calorimetry system (OxyCon Mobile, Vyaire Medical, Mettawa, IL) and applying standard indirect calorimetry equations^92^. To obtain net metabolic power for each condition, we subtracted the average metabolic power collected during the standing trials from metabolic power recorded during the walking conditions. To estimate steady-state, mass-specific, net rate of metabolic energy expenditure, we averaged the breath-by-breath data for the last minute of each five-minute trial and divided by body mass.

### Joint kinematics and kinetics measurements

We measured lower-limb joint kinematics using a reflective marker motion capture system (120Hz, Vicon, Oxford, UK) whereby the participant was instrumented with 44 reflective markers to capture the 6-DOF motion of the foot, shank, thigh, and pelvis. We calculated joint angles from the marker data and joint velocity was calculated as the first derivative of joint angle (Visual 3D, C-Motion Germantown, MD). A split-belt instrumented treadmill (980Hz, Bertec, Columbus, OH) measured ground reaction forces (GRFs). We performed inverse dynamic analysis to calculate net joint moments (Nm kg^-1^) for the ankle, knee, and hip (Visual 3D, C-Motion Germantown, MD). We filtered marker positions at 6Hz and analog data (*e.g.*, GRFs, exoskeleton forces) were filtered at 25Hz. We calculated biological contribution to ankle moment by subtracting the directly measured exoskeleton torque from the net ankle moment. Joint angles and moments are reported for the sagittal plane. We calculated rigid foot, 6-DOF joint power using techniques similar to Zelik *et al* ^93^.

For each participant, we calculated an average stride for each of the 5 conditions by time normalizing each stride between heel-strike and heel-strike of the subsequent stride. We then averaged multiple strides (15.5 ± 1.1) together to obtain a single normalized stride for each condition. We calculated integrated and peak values prior to inter-stride averaging. Peak values for a given measurement and condition were the average of the peaks for each stride within that condition. Stride and stance average mass-specific joint moment and power was the integral of the moment/power for each stride/stance period and dividing by stride/stance time and participant mass. To obtain the average moment per unit time, we divided again by the stride/stance time over which the integral was taken to obtain the average moment rate (Nm kg^-1^ s^-1^). (Note: This rate represents an average moment per unit time and is not a measure of how rapidly the moment is generated)

### Muscle activity measurements

We measured muscle activity of the ankle plantar flexors (medial and lateral gastrocnemius, soleus) and ankle dorsiflexor (tibialis anterior) using surface electromyography on the left leg (SX230, Biometrics Ltd, Newport, UK). To obtain an EMG envelope, we high-pass filtered at 20Hz the raw EMG data, rectified, and low-pass filtered at 10Hz (zero-phase 2^nd^ order Butterworth). Integrated EMG (iEMG) was the time-integral of the EMG envelope averaged across each stride for a condition. We then normalized amplitude of the EMG envelope and the iEMG for each muscle to the peak amplitude for the muscle across all conditions for each participant. Then, we divided the iEMG by the time period over which it was integrated to get the average EMG. Finally, to get average muscle activation per unit time, we again divided the average EMG by the time period over which it was averaged to obtain average muscle activation (EMG) rate (s^-1^). Average muscle activation rates over distinct phases of the stride was the average muscle activation over that phase divided by the time accrued during that phase. Phases examined were: all of stance (0-60%), early stance (0-40%) and late stance (40-60%) of the stride. (Note: This rate represents an average activation per unit time and is not a measure of how rapidly the muscle turns on and off.)

### Muscle-tendon dynamics measurements

We recorded B-mode ultrasound images of the soleus muscle using a linear probe ultrasound system (LV 7.5/60/96Z, Telemed, Lithuania). We positioned the probe over the right soleus below the contact point with the exoskeleton and secured with elastic adhesive wrap. We digitized ultrasound images using automated tracking software to determine time-varying soleus fascicle lengths and pennation angles during walking^94-96^. We manually selected a region of interest (soleus muscle belly) and the fascicle of interest, in the mid-region of the muscle belly, in an initial frame. The algorithm then tracks the fascicle in sequential frames by implementing an affine flow model. The UltraTrack^96^ program also incorporates a ‘key-frame correction’ algorithm that assists with removing temporal drift. For each step, we determined the corresponding index for the maximum dorsiflexion angle (from synced motion capture) and used it in the key-frame correction. However, key-frame correction assumes that errors accumulate linearly, which is not always appropriate for cyclical movements. When key-frame correction did not provide satisfactory results and tracking results were poor, manual corrections were made. Manual corrections were most commonly made during the swing phase, when fascicle lengthening was greatest. A square wave digital pulse from the ultrasound system initiated the capture synched the US data with the motion capture data. We incorporated the timing delay between systems in the analysis.

We calculated plantar flexor muscle-tendon (MT) force and length from biological ankle joint kinematics and kinetics^97^ using methods similar to previous studies ^69^. We calculated total MT force *F*_*MT*_, by dividing the net ankle joint moment *M*_*ank*_, by the MT moment arm about the ankle joint center *ma*_*ank*_, over the stride. The moment arm was derived from the relationship between the MTU length and the measured ankle angle similar to previous work^69^. Then, we estimated the contribution from the soleus to total MT force as the ratio of the cross-sectional area (CSA) of the soleus muscle to the total plantar flexor muscle cross sectional area (=0.54)^72^. Next, we calculated soleus muscle force 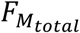, by dividing the soleus MT force contribution by the cosine of the measured soleus fascicle pennation angle *θ*_*p*_, over the stride (Eq. 2; equivalent to Eq. S3).

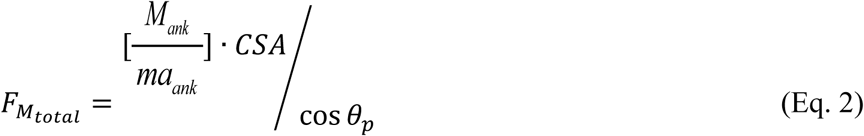

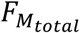 was divided by participant mass to get a mass normalized soleus muscle force (N kg^-1^). Soleus muscle fascicle velocity was the first derivative of fascicle length. For each participant, we calculated time-domain measurements for each condition by time normalizing each stride between heel-strike and heel-strike of the subsequent stride. We averaged together a minimum of three normalized strides (6 ± 0.6 stride average) to obtain a single normalized stride for a given condition. Integrated and peak values were calculated prior to inter-stride averaging. We calculated soleus muscle fascicle length at the time of peak force with the intention of observing changes in fascicle length due to changes in MT stretch. For each condition, we then averaged the length measured at peak force for at least three strides. Researchers were not blinded to the conditions during processing, but conditions were processed in random order.

Finally, to evaluate the influence of force-length (F-L) and force-velocity (F-V) effects on the capacity of the soleus muscle to produce force we computed its force per activation (N kg^-1^) (see S1, Eq. S7 for details on this metric). To do this we divided the average muscle force rate by the average muscle activation rate over the same distinct phases of the stride described previously for force and activation measures (*i.e.*, all of stance (0-60%), early stance (0-40%) and late stance (40-60%) of the stride).

### Statistics

Statistical analyses were performed on the exoskeleton conditions (kexo= 0-250 Nm rad^-1^) to isolate the role of exoskeleton rotational stiffness rather than the effect of the added mass of the exoskeleton and other structural impacts of the device. For ankle joint dynamics (n=11), muscle-tendon dynamics (n=10), muscle activity (n=10), and net metabolic energy rate (n=11), we report the means and standard errors calculated across participants. We note that dynamic ultrasound data was excluded for 1 of the 11 participants due to inability to track that participant’s data. Muscle level analysis for this participant was excluded from our analyses. Metrics describing soleus muscle fascicle dynamics (*i.e.*, force, activation, length, velocity) were the primary outcome measures examined in this study. To address our hypotheses, we performed a series of two-factor ANOVA (mixed-model, random effect: participant; main effect: k_exo_) analyses to test for an effect of exoskeleton stiffness k_exo_, on soleus muscle force rate, activation rate, fascicle length, fascicle velocity, and force per activation (JMP Pro, SAS, Cary, NC). A Shapiro-Wilk W test confirmed normality of the data. For dependent variables showing a significant main effect (*p< α*=0.05) of exoskeleton stiffness, a Tukey’s Honestly Significant Difference Test (THSD)^98^ was used to reveal pairwise differences between conditions. Finally, we performed a number of within-participant, linear, least-squares regression (LLSR) analyses to test for relationships (*p*< α = 0.05; R^2^) between changes in soleus muscle neuromechanics (*e.g.*, positive power = positive work rate, force, force rate, activation, activation rate, force /activation) and changes in whole body net metabolic rate due to increasing exoskeleton stiffness.

### Data availability

Source data from this study in .mat and .txt format and an associated readme.txt for navigating it are available for download at: http://pwp.gatech.edu/hpl/archival-data-from-publications/

## Supporting information

Supplemental Analysis

## Additional Information

## Acknowledgments

We thank S. Steele-Pardue and T. Giest for assistance in assembling the exoskeleton device, T. Giest, J. McCall, and S. Philius for help with data collection and F. Shaw for help with data analysis.

## Author contributions

R.W.N. and G.S.S. contributed equally to study design and direction; R.W.N. designed and fabricated the exoskeleton device; R.W.N conducted the human locomotion experiments; R.W.N., T.J.D, O.N.B. and G.S.S. analyzed the data; R.W.N., T.J.D., O.N.B., and G.S.S. drafted the manuscript; R.W.N., T.J.D., O.N.B. and G.S.S. edited the manuscript. All authors approved the final manuscript.

