## Supplemental Analysis for "Ultrasound imaging links soleus muscle neuromechanics and energetics during human walking with elastic ankle exoskeletons"

**Title**

Richard W. Nuckols
Taylor J.M. Dick
Owen N. Beck
Gregory S. Sawicki

**Influence of increasing elastic ankle exoskeleton stiffness on the metabolic cost of soleus muscle force production**

The metabolic energy utilized by leg muscles to generate ground force is a primary determinant of metabolic rate during locomotion[1-10]. In fact, the relationship between metabolic rate and the rate of muscle force production during human walking has been formalized by Griffin et al[8] and is recapitulated here in a slightly different form to focus on the soleus muscle during stance phase (Eq. S1 below); rather than *all* of the leg extensor muscles during ground contact (Eq. 3 in Griffin et al[8]). Namely, we assume that the soleus muscle consumes metabolic energy at a rate ($\dot{E}_{met}$) that is proportional to its rate ($\frac{1}{T_{act}}$) of active muscle volume ($V_{act}$); the volume of active muscle used to generate force per unit activation time ($\frac{V_{act}}{T_{act}}$) (Eq. S1)

$$\dot{E}_{met} = \alpha \cdot \frac{V_{act}}{T_{act}} \qquad \text{(Eq. S1)}$$

The constant '$\alpha$' is the amount of soleus metabolic energy consumed per unit muscle volume (J $cm^{-3}$). Consistent with previous research[8], we assume that changes in the mechanical demand of the task, in this case due to changing ankle exoskeleton stiffness, does not affect $\alpha$, and that soleus is active during *all* of stance phase (*i.e.*, $T_{act} = T_{stance}$). This simple equation (Eq. S1) provides a framework to examine how changes in soleus muscle dynamics and activation during walking with elastic ankle exoskeletons impact soleus metabolic energy consumption by asking: How do elastic ankle exoskeletons influence the rate of active soleus muscle volume?

We start by defining active muscle volume ($V_{act}$) as the product of a muscle's activation (*act*), physiological cross-sectional area (*PCSA*), and fascicle optimal length ($l_0$) (Eq. S2).

$$V_{act} = act \cdot PCSA \cdot l_0 \qquad \text{(Eq. S2)}$$

While there is some evidence that $l_0$ may change with muscle activation [11,12], here we assume that soleus $l_0$ and *PSCA* are constants that do not change with increasing elastic ankle exoskeleton stiffness. As a result, active muscle volume ($V_{act}$) is directly proportional to muscle activation (*act*) across all exoskeleton stiffness conditions (Eq. S2). It follows then, from Eq. S1 that changes in the rate of $V_{act}$ ought to correlate with changes in metabolic rate of the soleus and by extension, reflect changes in whole-body metabolic rate during walking with elastic ankle exoskeletons over a range of stiffness conditions. Indeed, our electromyography measurements, which serve as proxy for muscle activation

($act$), indicated a moderate correlation ($R^2$=0.44) between change in stance phase soleus activation rate (see Methods for details on rate calculations) and changes in whole-body net metabolic rate with respect to the zero exoskeleton stiffness condition (Table 2). Our finding that changes in muscle activation well-explains  metabolic rate during walking with exoskeletons is supported by previous work using powered ankle exoskeletons (see Fig. 9, Jackson et al.[13]). Furthermore, if the correlation is causal, a linear fit to the Δ net metabolic rate (W $kg^{-1}$) versus Δ soleus activation rate (unitless $s^{-1}$) data suggest that for this exoskeleton a unit reduction in soleus activation rate should yield a ~1.3 W $kg^{-1}$reduction in whole-body net metabolic rate (Fig. 5). In theory this was achieved by reducing $V_{act}$ and/or increasing $T_{stance}$.

To gain further insight into how elastic ankle exoskeletons reduce soleus $V_{act}$ rate over the stance phase of walking we took a top-down approach to examine how applying exoskeleton torque to changes in underlying soleus muscle activation ($act$). We start at the joint-level by estimating the total soleus fascicle force ($F_{M_{total}}$), from the force on the combined plantar flexor muscle-tendons ($F_{MT}$) computed via inverse dynamics, accounting for the soleus's relative cross-sectional area (CSA) within the triceps surae group (CSA=0.56, see Methods Eq. 2) as well as its instantaneous pennation angle ($\theta_p$) (Eq. S3).

$$F_{M_{total}} = {F_{MT} \cdot CSA}\Big/{\cos\theta_p} \qquad \text{(Eq. S3)}$$

Elastic ankle exoskeletons directly reduce the force on the plantar flexor ($F_{MT}$) because they contribute torque in parallel with the ankle-joint that offsets the biological moment and mechanical power required to walk (Fig. 2C, D)[14,15]. In addition, our data indicate that increasing exoskeleton stiffness does not alter muscle pennation angle (Supp Fig. 3D) and we assume that soleus *CSA* is constant and independent of exoskeleton stiffness. Thus, according to Eq. S3, reductions in plantar flexor MT force ($F_{MT}$) due to elastic ankle exoskeleton torque assistance should directly translate to reductions in soleus total force $F_{M_{total}}$.

When linking muscle total force ($F_{M_{total}}$) and its activation ($act$ ~=$V_{act}$), it is important to consider whether all of the force is generated by active contractile machinery. For example, it is well known that when a muscle is stretched beyond its optimal length ($l_0$) some of the total muscle force ($F_{M_{total}}$) is contributed by structures in parallel with the contractile apparatus that can produce muscle force passively ($F_{M_{pas}}$) (Fig. 4B, dashed curve). Thus, for relatively long muscle operating lengths, active muscle force ($F_{M_{act}}$) requirements could be reduced by passive contributions (Eq. S4).

$$F_{M_{act}} = F_{M_{total}} - F_{M_{pas}} \quad \text{(Eq. S4)}$$

Even for walking with the stiffest exoskeletons we tested ($k_{exo}$=250 Nm rad$^{-1}$), we estimate that passive muscle force was likely not a factor (Fig. 4C). We note however, that for elastic ankle exoskeletons stiffer than 250 Nm rad$^{-1}$, passive forces may contribute due to the trend toward increasing soleus operating lengths (Fig. 4A). Given that in this study active and total soleus muscle force were near equivalent ($F_{M_{act}} \cong F_{M_{total}}$) (Eq. S4), any reduction in $F_{M_{act}}$ due to the exoskeleton should be captured by changes in $F_{M_{total}}$. Our analysis indicated a moderate correlation ($R^2$=0.42) between changes in soleus total force rate during early stance (Fig. 3A) (see Methods for details on rate calculations) and changes in whole-body net metabolic rate with respect to the zero exoskeleton stiffness condition (Table 2).

It is tempting to stop here and conclude that elastic ankle exoskeletons reduce metabolic rate merely by reducing soleus active muscle force rate during early stance, which by itself reduces soleus activation rate over stance, and ultimately reduces the volume of active muscle per unit activation time ($\frac{V_{act}}{T_{act}}$) (Eq. S1). But, the Hill-type model of muscle force production[16] highlights that force is not equivalent to activation and that other physiological parameters (F-L, F-V) must be considered. Using a Hill-type model, we estimate a muscle's active force output as the product of maximum muscle force ($F_{M_{max}}$), normalized activation (*act*) (defined as between 0 (not active) and 1 (fully active)) as well as dimensionless factors due to the force-length (F-L) (Fig. 4C) and force-velocity (F-V) (Fig. 4F) relationships (Eq. S5).

$$F_{M_{act}} = F_{M_{max}} \cdot act \cdot (F\text{-}L) \cdot (F\text{-}V) \quad \text{(Eq. S5)}$$

Solving Equation S5 for muscle activation, *act* yields:

$$act \cong \frac{F_{M_{act}}}{F_{M_{max}}} \cdot \frac{1}{(F\text{-}L) \cdot (F\text{-}V)} \quad \text{(Eq. S6)}$$

Since muscle force does not always reflect muscle activation (Eqs. S4-S6)[17], in addition to reducing soleus force rate, elastic ankle exoskeletons may also act to reduce activation by improving the economy of muscle force production. Combining Eq. 4 and 6, we see that, independent of the muscle force required, exoskeletons could shift soleus' operating point on the F-L and/or F-V curve to a point with greater force capacity (*e.g.*, up to or beyond the optimal length, *lo* or closer to isometric) yielding improved economy by increasing the muscle force per unit muscle activation (Eq. S7) (*i.e.*, by decreasing the activation and thus metabolic energy consumed per unit force [18-21]).

$$\frac{F_{M_{tot}}}{act} = F_{M_{max}} \cdot (FL) \cdot (FV) \cdot (1 + \frac{F_{M_{pas}}}{F_{M_{act}}}) \quad \text{(Eq. S7)}$$

Indeed, our data indicate that during early stance, increasing exoskeleton stiffness increased soleus operating length, pushing it toward its optimal length, $l_0$ (Fig. 4C), and increasing its force per activation (Fig. 4B). However, increasing exoskeleton stiffness also increased soleus shortening velocity in late stance pushing it toward $v_{max}$ (Fig. 4F) and decreasing its force per activation (Fig. 4E).

It is difficult to disentangle the effects of increasing ankle exoskeleton stiffness on soleus F-L and F-V effects as they relate to metabolic rate, because they tend to counteract each other's effects. This trade-off is perhaps reflected in the lack of correlation between changes in force per activation during stance and changes in whole-body net metabolic rate (Table 2). Nevertheless, our data suggest that the F-V driven decrease in soleus force per activation in late stance (Fig. 4E,F) outweighed the F-L driven increase in soleus force per activation in early stance (Fig. 4B,C) and contributed to the observed increase in both stance phase soleus muscle activation rate (Fig. 3B) and whole-body net metabolic rate (Fig. 2A) for the stiffest ankle exoskeletons. This notion is supported by a moderate negative correlation ($R^2$=0.53) between changes in soleus force rate in late stance (Fig. 3A) and changes in whole-body net metabolic rate (Fig. 2A) with respect to the zero exoskeleton stiffness condition (Table 2). That is, the stiffest exoskeletons, which also yielded the highest whole-body net metabolic rates were associated with reduced soleus muscle force rates in late stance (Fig. 3A); a consequence that is likely derived from poor F-V contractile conditions (Fig. 4F). As a result, in late stance, markedly reduced soleus force rate (Fig. 3A) did not translate to reductions in soleus muscle activation rate (Fig. 3B). This suggests that for the stiffest exoskeletons, reduced force capacity (*i.e.*, force per activation) (Eq. S7) (Figs. 4B,E) outweighed reduced force demand (Eqs. S3-S4) (Fig. 3A) and increased whole-body net metabolic rate (Fig. 2A). Taken together, these data support our hypothesis that a trade-off between reduced soleus force demand and a shift to soleus fascicle contractile conditions that are less economical for force production (*i.e.*, reduced force capacity) , both contribute to an intermediate stiffness 'sweet-spot' where users can use elastic ankle exoskeletons to reduce their metabolic rate during walking.
