## Supplemental Figure for "Ultrasound imaging links soleus muscle neuromechanics and energetics during human walking with elastic ankle exoskeletons"

**Title**

Richard W. Nuckols
Taylor J.M. Dick
Owen N. Beck
Gregory S. Sawicki

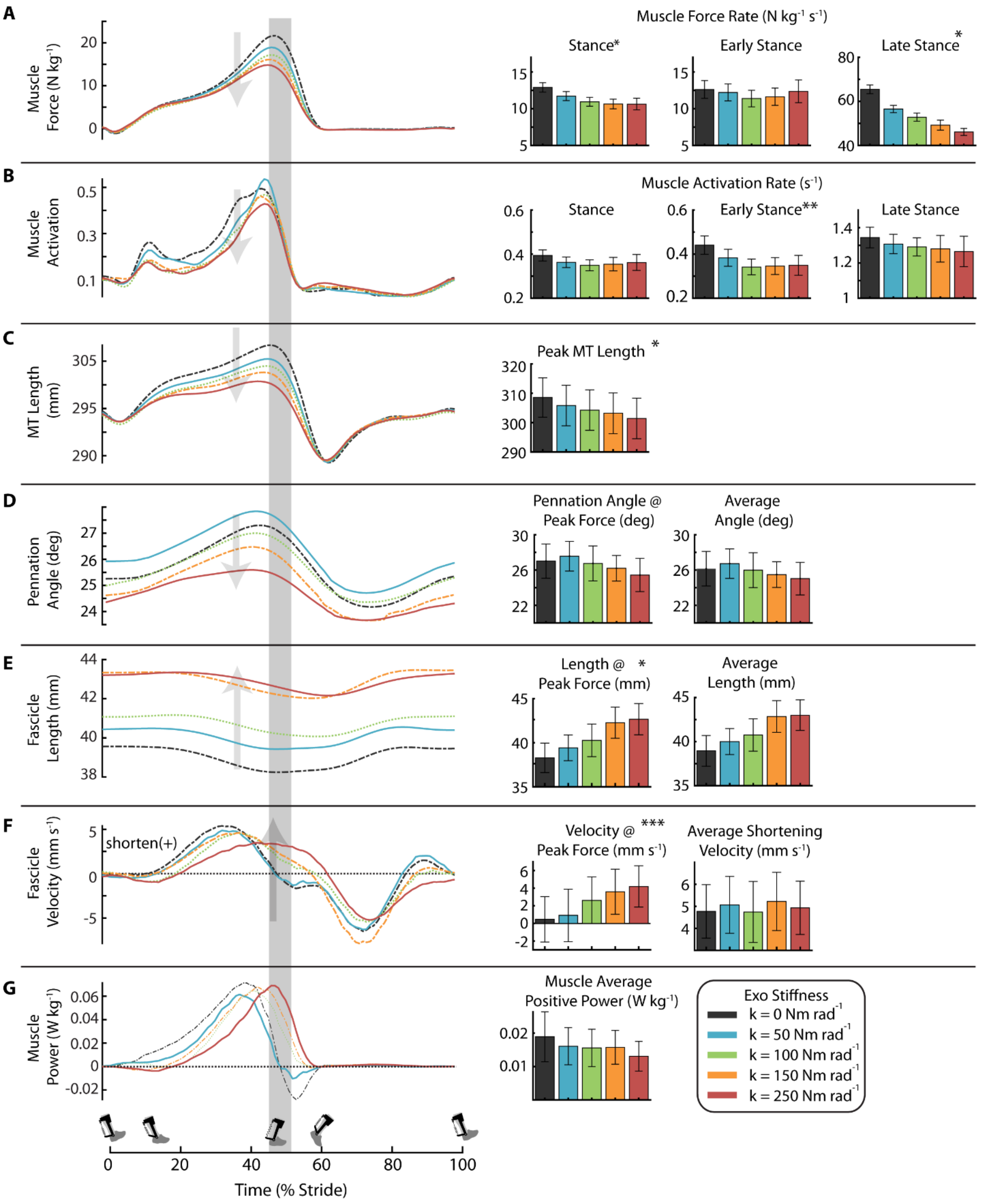
A
Muscle Force (N kg-1)
Muscle Force Rate (N kg-1 s-1)
Stance*
Early Stance
Late Stance*
B
Muscle Activation
Muscle Activation Rate (s-1)
Stance
Early Stance**
Late Stance
C
MT Length (mm)
Peak MT Length *
D
Pennation Angle (deg)
Pennation Angle @ Peak Force (deg)
Average Angle (deg)
E
Fascicle Length (mm)
Length @ * Peak Force (mm)
Average Length (mm)
F
Fascicle Velocity (mm s-1)
shorten(+)
Velocity @ *** Peak Force (mm s-1)
Average Shortening Velocity (mm s-1)
G
Muscle Power (W kg-1)
Muscle Average Positive Power (W kg-1)
Exo Stiffness
k = 0 Nm rad-1
k = 50 Nm rad-1
k = 100 Nm rad-1
k = 150 Nm rad-1
k = 250 Nm rad-1
Time (% Stride)

**Supplementary Figure 1: Effect of elastic ankle exoskeleton stiffness on soleus muscle-tendon dynamics. (A)** Soleus force for each exoskeleton stiffness averaged over participants. Region of peak force highlighted in time series data. Bar charts represent the soleus muscle force rate for stance, early stance (0-40% stride), and late stance (40-60% stride). Increasing exoskeleton stiffness resulted in decreased soleus force rate for stance and late stance. **(B)** Time series of soleus muscle activation where amplitude is normalized to the peak activation across all stiffnesses for each participant. Bar charts are soleus muscle activation rate in stance, early stance, and late stance. Soleus activation rate decreased with increasing exoskeleton assistance in early stance. **(C)** Time series of muscle-tendon unit length and bar charts representing the maximum MT length. Peak MT length decreased with increasing stiffness. **(D)** Time series of soleus fascicle pennation angle over stride. Bar charts represent the pennation angle at peak force and average angle during stance. Pennation angle trends towards a decrease angle with increasing stiffness though not significantly. **(E)** Time series of soleus fascicle length and bar charts representing length at peak fascicle force and average length during stance. Fascicle length increased with increasing exoskeleton stiffness. **(F)** Time series of soleus fascicle velocity measured and bar charts representing velocity at peak fascicle force and average shortening velocity during stance. Fascicle shortening velocity at peak increases with stiffness. **(G)** Time series of soleus fascicle power and bar charts representing muscle average positive power. All bars graphs show mean ± SEM across N=10 participants. [main effect: stiffness, * $p < 0.0001$, **$p = 0.0023$, ***$p = 0.0318$].
